# MAPK Pathway Inhibition as A Rational Therapeutic Strategy for MiR-138-5p/PAQR3 Dysregulation-mediated Epirubicin Resistance in Triple-negative Breast Cancer

**DOI:** 10.1101/2021.02.28.433242

**Authors:** Jianbo Huang, Shiyan Zeng, Yun Xiao, Xiaoyi Wang, Hongyuan Li, Tingxiu Xiang, Lingquan Kong, Guosheng Ren

**Author notes:** Correspondence should be addressed to G. R., Department of Endocrine and Breast Surgery, The First Affiliated Hospital of Chongqing Medical University, Youyi Road 1, Chongqing, 400016, China.

## Abstract

Anthracyclines, such as epirubicin, activate the mitogen-activated protein kinase (MAPK) pathway in breast cancer (BC). Better predictors of tumor response are needed to guide de-intensification of anthracyclines when used in BC treatment. Here, we aimed to see if MAPK activation was responsible and targetable for epirubicin resistance, and to explore mechanisms and predictive markers for resistance. MAPK pathway inhibitors were used to ameliorate epirubicin resistance. Negative regulators of MAPK were screened to identify the essential gene cascades required for activation of the pathway. In vitro and in vivo approaches were applied to investigate epirubicin resistance. The regulatory miRNA was identified through bioinformatics-based screening and luciferase reporter assay. 114 estrogen receptor negative patients who received epirubicin monotherapy were included to evaluate predictive markers for tumor response. The MAPK pathway was activated in cells resistant to epirubicin; MAPK inhibition ameliorated epirubicin resistance. In triple-negative breast cancer (TNBC), progestin and adipoQ receptor 3 (PAQR3) were the most significantly decreased negative MAPK regulators. PAQR3 increased epirubicin sensitivity and suppressed MAPK activation in resistant cells. MiR-138-5p was increased in epirubicin resistant cells, and was shown to downregulate PAQR3, causing resistance in epirubicin resistant cells. PAQR3 was an independent factor in predicting response to epirubicin (OR=4.86, 95%CI=1.13-20.87, P=0.034), and possessed a high negative predictive value (NPV) (0.93; 95%CI=0.83-0.97). In addition, PAQR3 exhibited a better predictive value in patients older than 50 years (12.92 (95% CI, 1.43-116.78, P = 0.023). The combinative use of PAQR3 and topoisomerase II-α (TOP2A) led to an increased specificity (0.70; 95%CI=0.61-0.79), when compared with either PAQR3 or TOP2A alone. MAPK inhibition ameliorated epirubicin resistance. MiR-138-5p-induced PAQR3 reduction causes epirubicin resistance in TNBC via activation of MAPK cascades. PAQR3 is an independent and favorable predictor response to epirubicin in BC.

## Introduction

Breast cancer (BC), the most common malignancy for global females, has been arising for decades. It is estimated accounting for 30% of the new cases of all sites in 2019. Although the average 5-year survival rate of all stages and races elevates to 90%, BC remains a leading cause of cancer death in less developed countries and the second leading cause of cancer death in American women, exceeded only by lung cancer ^[1]^.

Anthracyclines, the efficient cancer cell killer through poisoning of the nuclear enzyme topoisomerase II, give abundant benefits to both early- and advanced-stage BC, nevertheless, resistance to anthracyclines is inevitable in fractional tumor cells. So, unmasking the underlying mechanisms would shed much light on BC novel treatment development. Intriguingly, previous evidences showed anthracyclines activated p44/42-mitogen-activated protein kinase (MAPK) pathway, which elevates cell death threshold then facilitates surviving in cancer cells ^[2, 3]^.

In this study, we showed MAPK pathway was activated in epirubicin-resistant BC cells and was targetable. Different from in other subtypes of BC, a non-coding RNA-based regulation was recovered to elucidate MAPK activation-mediated epirubicin resistance in triple-negative breast cancer (TNBC). Meaningfully, we proved progestin and adipoQ receptor 3 (PAQR3), a recently identified Raf kinase trapper, was more preferable than topoisomerase II alpha (TOP2A) expression in predicting the response to anthracyclines-based chemotherapy.

## Results

### MAPK pathway activation was responsible and targetable for epirubicin resistance

The half maximal inhibitory concentration (IC_50_) obviously increased (Fig. 1A). The resistant cells exhibited strengthened survivability (Fig. 1B), and showed an average 2.0-, 1.5- and 1.9-fold increase in colony formation ability (Fig. 1C).

**Figure 1.**
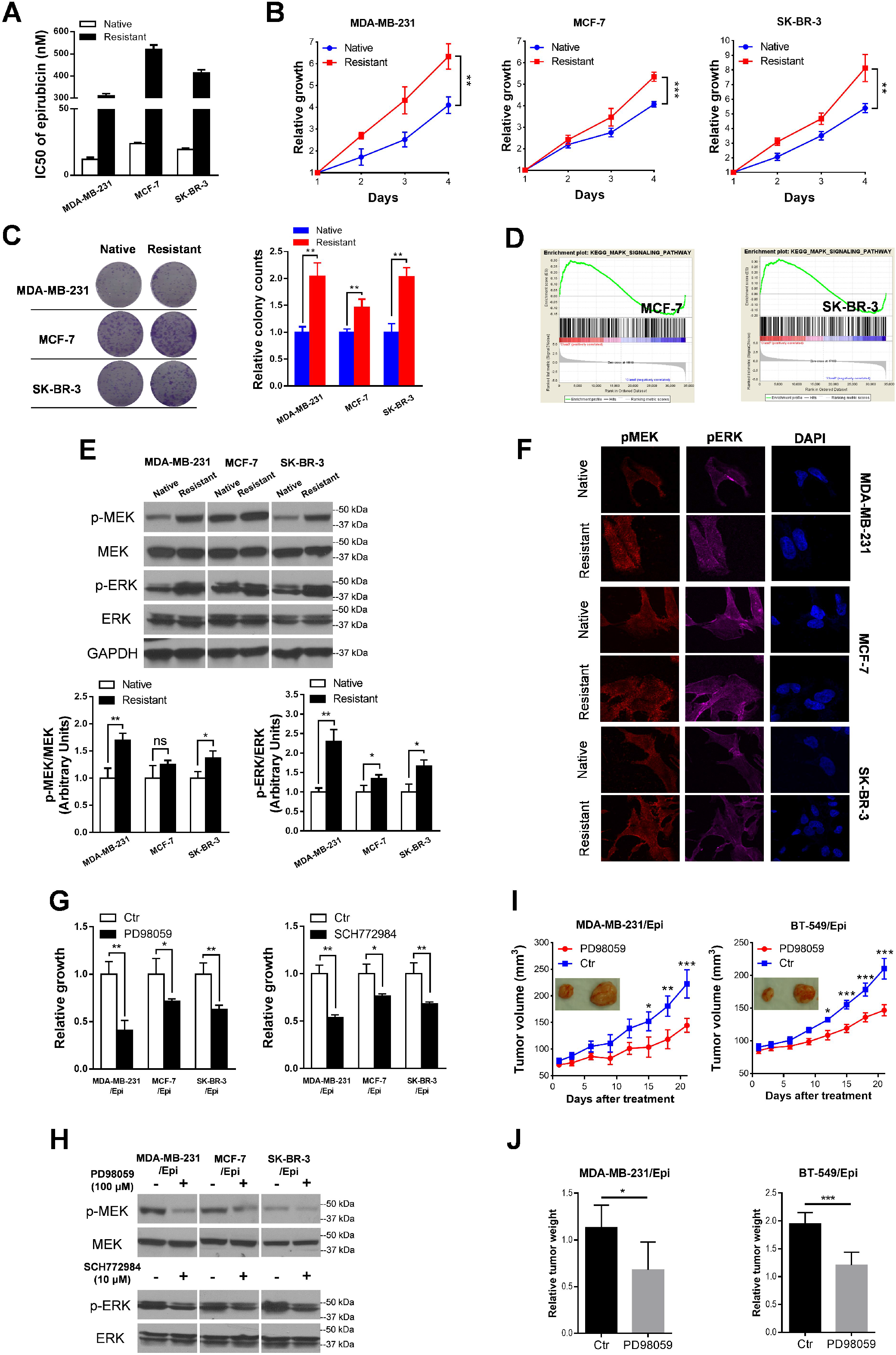
Inhibiting MAPK pathway activation ameliorated epirubicin resistance in breast cancer cells. (A) The half maximal inhibitory concentration (IC_50_) obviously increased in MDA-MB-231/Epi (from 12.1 to 311.3 nM), MCF-7/Epi (from 23.8 to 520.8 nM), and SK-BR-3/Epi cells (from 19.4 to 415.1 nM). (B) The epirubicin-resistant breast cancer cells were characterized with an increased cell viability as compared to pararenal native cells, after incubation with 0.5 μM epirubicin for 24, 48 and 72 hours. (C) MDA-MB-231/Epi, MCF-7/Epi and SK-BR-3/Epi cells showed an average 2.0-, 1.5- and 1.9-fold increase in colony formation ability, respectively, after 0.3 μM epirubicin treatment for two weeks. (D) Gene set enrichment analysis (GSEA) showed enrichment of MAPK pathway components after epirubicin resistance induction in MCF-7 and SK-BR-3 cells. Western blot (E) and immunofluorescence (F) assays showed an obviously increased levels of phosphorylated MEK1 and ERK1/2 in resistant cells when compared with native cells. (G) Either MEK1/2 (PD98059) or ERK1/2 inhibitor (SCH772984) was able to suppress epirubicin-resistant cells viability. (H) The levels of phosphorylated MEK1/2 and ERK1/2 were obviously ameliorated after inhibitors addition. The in vivo study showed MEK inhibitor PD98059 reduced tumor volume (I) and weight (J) in MDA-MB-231/Epi and BT-549/Epi cells. ns: no significance, * *P* < 0.05, ** *P* < 0.01, *** *P* < 0.001.

Epirubicin potentiated ERK1/2 phosphorylation in BC cells ^[3]^, here we aimed to study if epirubicin-induced ERK1/2 phosphorylation was responsible for BC epirubicin resistance. Similar to previous finding in MDA-MB-231/Epi cells ^[4]^, MAPK signaling was more inclined to be activated in resistant cells (MCF-7/Epi and SK-BR-3/Epi cells) through analyzing the transcriptome dataset (Fig. 1D). Consistently, phosphor-MEK1/2 and ERK1/2 levels were increased in epirubicin-resistant cells when compared with native cells (Fig. 1E and F). Summary, these results suggested MAPK pathway activation in epirubicin-resistant BC cells. Subsequently, to prove activated MAPK pathway was responsible and targetable for epirubicin resistance, MEK1/2 (PD98059) and ERK1/2 inhibitors (SCH772984) were added to treat resistant cells. After 96 hours incubation, PD98059 suppressed cell viability by 59.2% for MDA-MB-231/Epi cells, 28.6% for MCF-7/Epi cells and 37.3% for SK-BR-3/Epi cells, SCH772984 inhibited cell proliferation by 46.1% for MDA-MB-231/Epi cells, 23.5% for MCF-7/Epi cells and 31.7% for SK-BR-3/Epi cells (Fig. 1G). Consistently, the reduced levels of phosphorylated MEK1/2 and ERK1/2 were observed after inhibitors treatment (Fig. 1H). The in vivo study showed MEK inhibitor PD98059 reduced tumor volume and weight in MDA-MB-231/Epi and BT-549/Epi cells (Fig. 1I and J).

### PAQR3 expression significantly decreased in epirubicin-resistant TNBC cells

To elucidate mechanisms responsible for MAPK pathway activation in epirubicin-resistant BC cells, we hired a literature-based gene signature named negative regulation of MAPK pathway (Reactome, R-HSA-5675221) to screen the main candidates responsible for MAPK pathway activation within GSE54326 dataset. Interestingly, in MDA-MB-231/Epi cells, the most obviously altered MAPK pathway negative regulator was PAQR3, a recently identified ERK1/2 phosphorylation inhibitor through trapping Raf to Golgi apparatus (Fig. 2A). Real-time PCR assay further validated that PAQR3 had the most tremendous decrease in resistant cells (Fig. 2B), decrease of PAQR3 protein level was also confirmed (Fig. 2C). Similar to previous studies ^[5, 6]^, MEK1/2 and ERK1/2 phosphorylation were enhanced after downregulating PAQR3 expression, whereas levels of phosphor-MEK1/2 and ERK1/2 reduced with enforcing PAQR3 expression (Fig. 2D). In non-TNBC resistant cells, there was no obvious altered MAPK pathway regulator when compared with native cells (Fig. 2E and F).

**Figure 2.**
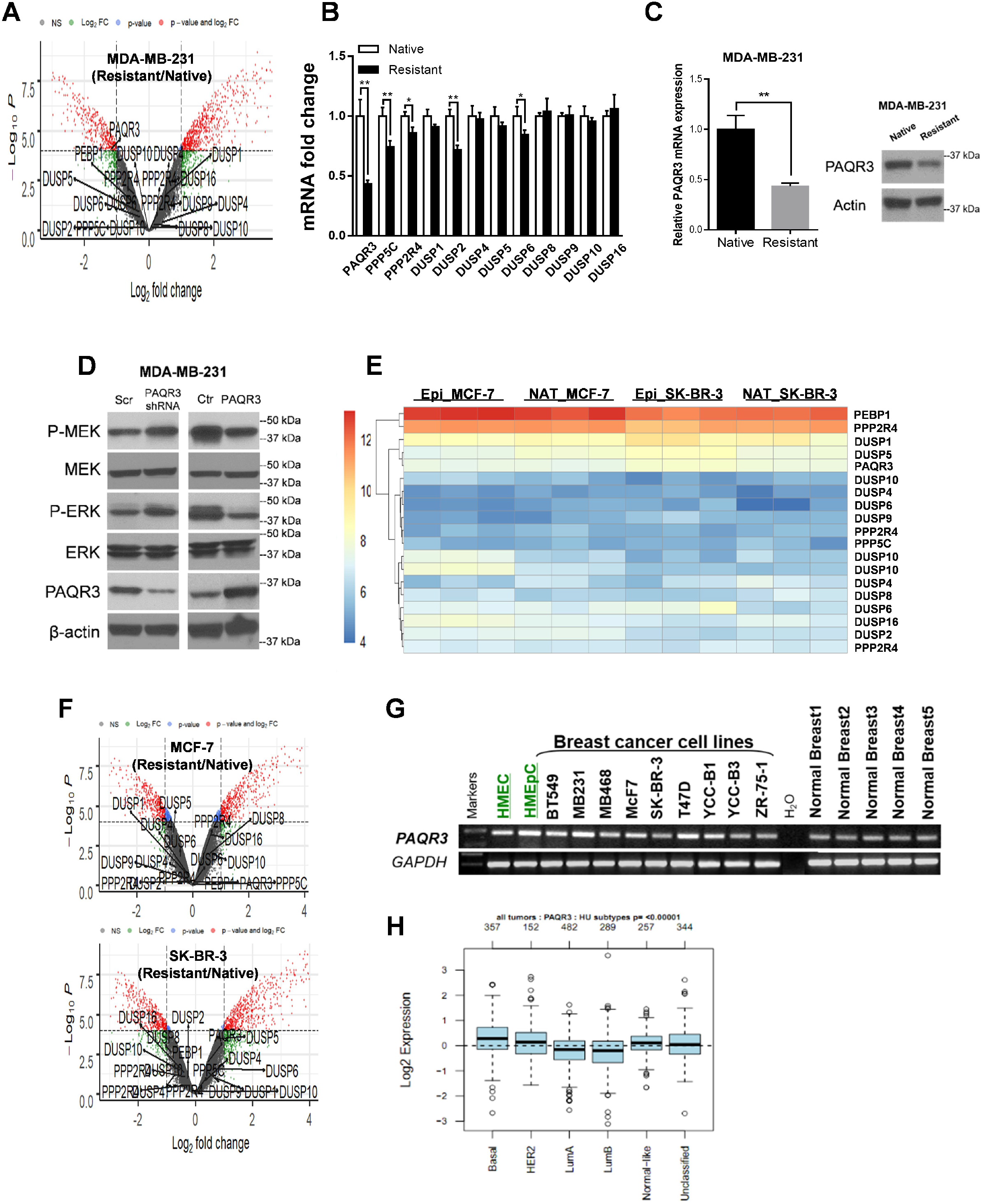
PAQR3 expression significantly decreased in MDA-MB-231/Epi cells. (A) In TNBC MDA-MB-231/Epi cells, PAQR3, the MAPK pathway negative regulator, most obviously decreased in epirubicin-resistant cells, whereas other well-known MAPK pathway negative regulators stayed unaltered. (B) Real-time PCR assay further validated that PAQR3 had the most tremendous decrease in epirubicin-resistant cells. (C) The decrease of PAQR3 protein level was confirmed with western blot assay. (D) Artificially suppressing PAQR3 expression by shRNA increased the phosphorylated levels of MEK1/2 and ERK1/2, whereas enforcing PAQR3 expression expectedly suppressed MEK1/2 and ERK1/2 phosphorylation. (E) The heatmap showed the expression of MAPK pathway negative regulators in epirubicin-resistant and native cells. (F) The volcano plots showed no regulator reached at the significance based on biological meaning (ļlog2fold changeļ>1, (ļlog10P valueļ>4) in MCF-7 and SK-BR-3 cells. (G) PAQR3 was broadly expressed in multiple subtypes of breast cancer cells. (H) The basal type was enriched with the highest PAQR3 expression ** *P* < 0.01.

PAQR3 was broadly expressed in multiple BC cell lines (Fig. 2G), and the basal type was enriched with the highest PAQR3 expression (Fig. 2H), together with the finding that PAQR3 most tremendously decreased in resistant TNBC, we inferred PAQR3 was vital for susceptibility to epirubicin cytotoxicity in TNBC.

### PAQR3 ameliorated epirubicin resistance in TNBC cells

We determined if PAQR3 reduction-caused ERK1/2 activation was responsible for epirubicin resistance. After knocking down the endogenous levels of PAQR3 (Fig. 3A), tumor cells became insensitive to epirubicin, with 1.8- and 2.6-fold increase in cell viability (Fig. 3B) and colony formation (Fig. 3D), respectively. The caspase 3/7 activity had a 62% decrease after downregulating PAQR3 (Fig. 3F). The increase in ERK1/2 and MEK1 activation was also observed (Fig. 3I). Inversely, resistant cell MDA-MB-231/Epi was used to re-express PAQR3 (Fig. 3A), followed with average 32% decrease in cell viability (Fig. 3C), 60% decrease in colony formation ability (Fig. 3E), and 2.8-fold increase in caspase3/7 activity (Fig. 3G). Also, the phosphorylation of MEK1/2 and ERK1/2 after PAQR3 re-expression obviously decreased in epirubicin-resistant cells (Fig. 3H and I).

**Figure 3.**
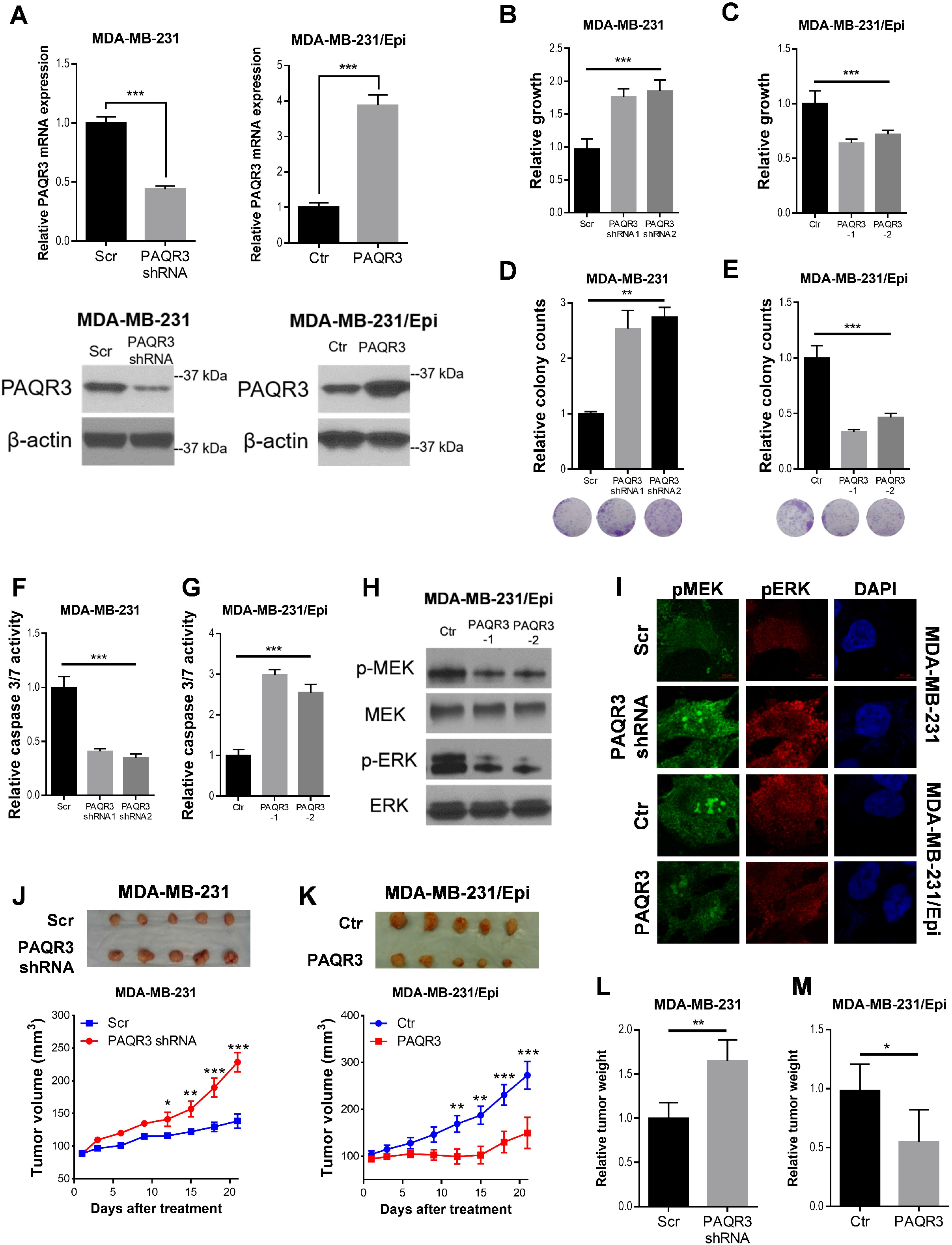
PAQR3 ameliorated epirubicin resistance through suppressing MAPK pathway activation in TNBC cells. (A) The endogenous PAQR3 levels in MDA-MB-231 cells were successfully suppressed with RNA interference technology. We also enforced PAQR3 expression in MDA-MB-231/Epi cells. Knocking down PAQR3 in native MDA-MB-231 cells induced 1.8- and 2.6-fold increase in cell viability (B) and colony formation ability (D), whereas enforcing PAQR3 expression in resistant cells ameliorated epirubicin resistance, characterized with average 32% decrease in cell viability (C) and 60% decrease in colony formation ability (E), as compared to control cells. (F) Knocking down the endogenous PAQR3 decreased caspase 3/7 activity by 62% after epirubicin incubation, (G) whereas enforcing PAQR3 expression induced a 2.8-fold increase in caspase3/7 activity. (H) PAQR3 inhibited phosphorylation of MEK1/2 and ERK1/2 in MDA-MB-231/Epi cells. (I) Immunofluorescence assay showed elevated phosphorylation of MEK1 and ERK1/2 after knocking down PAQR3 in MDA-MB-231 cells, and reduced levels of phosphorylated MEK1 and ERK1/2 in MDA-MB-231/Epi cells, following PAQR3 artificial enhancement The in vivo study showed native cells with PAQR3 suppressed exhibited a reduced sensitivity to epirubicin, featured with more aggressive tumor growth (J) and higher tumor weight (L). Inversely, enforcing PAQR3 in resistant cells enabled xenograft tumor more sensitive to epirubicin treatment, characterized with lower tumor volume (K) and weight (M). * *P* < 0.05, ** *P* < 0.01, *** *P* < 0.001.

We further characterized PAQR3’s role in epirubicin resistance by employing a subcutaneous xenograft model. Tumors with endogenous PAQR3 decrease presented 1.6-fold increase in tumor size (Fig. 3J) and weight (Fig. 3L), whereas tumors with high PAQR3 expression had less aggressive growth, with size decreased by 53% (Fig. 3K) and weight reduced by 44% (Fig. 3M).

### MiR-138-5p upregulation led to PAQR3 reduction in resistant TNBC cells

MiRNAs upregulation were involved in epirubicin resistance development of BC cells ^[7]^. Here, of the most upregulated miRNAs in MDA-MB-231/Epi cells (Fig. 4A), miR-138-5p was co-predicted targeting the 3’UTR of PAQR3 by three databases (Fig. 4B). Moreover, we observed 4.3- and 3.4-fold increase of miR-138-5p in resistant TNBC cells (Fig. 4C). The predicted interaction between miR-138-5p and targeting sites within the 3’-UTR of PAQR3 was illustrated (Fig. 4D). The luciferase reporter assay further proved PAQR3 was a direct target of miR-138-5p, as co-transfection of pre-miR-138-5p and luciferase reporter plasmid showed 47% decrease in luciferase activity. However, no significant difference of luciferase activity was observed in co-transfection of miRNA and mutant plasmid (Fig. 4E). Pre-miR-138-5p and anti-miR-138-5p were utilized to force and inhibit miR-138-5p expression (Fig. 4F). As expected, pre-miR-138-5p led to 48% and 38% decrease of PAQR3 mRNA in TNBC cells (Fig. 4G), whereas anti-miR-138-5p induced 2.0- and 1.8-fold increase in PAQR3 in resistant cells (Fig. 4H). Also, the PAQR3 protein levels correspondingly changed after regulating miR-138-5p expression (Fig. 4I and J).

**Figure 4.**
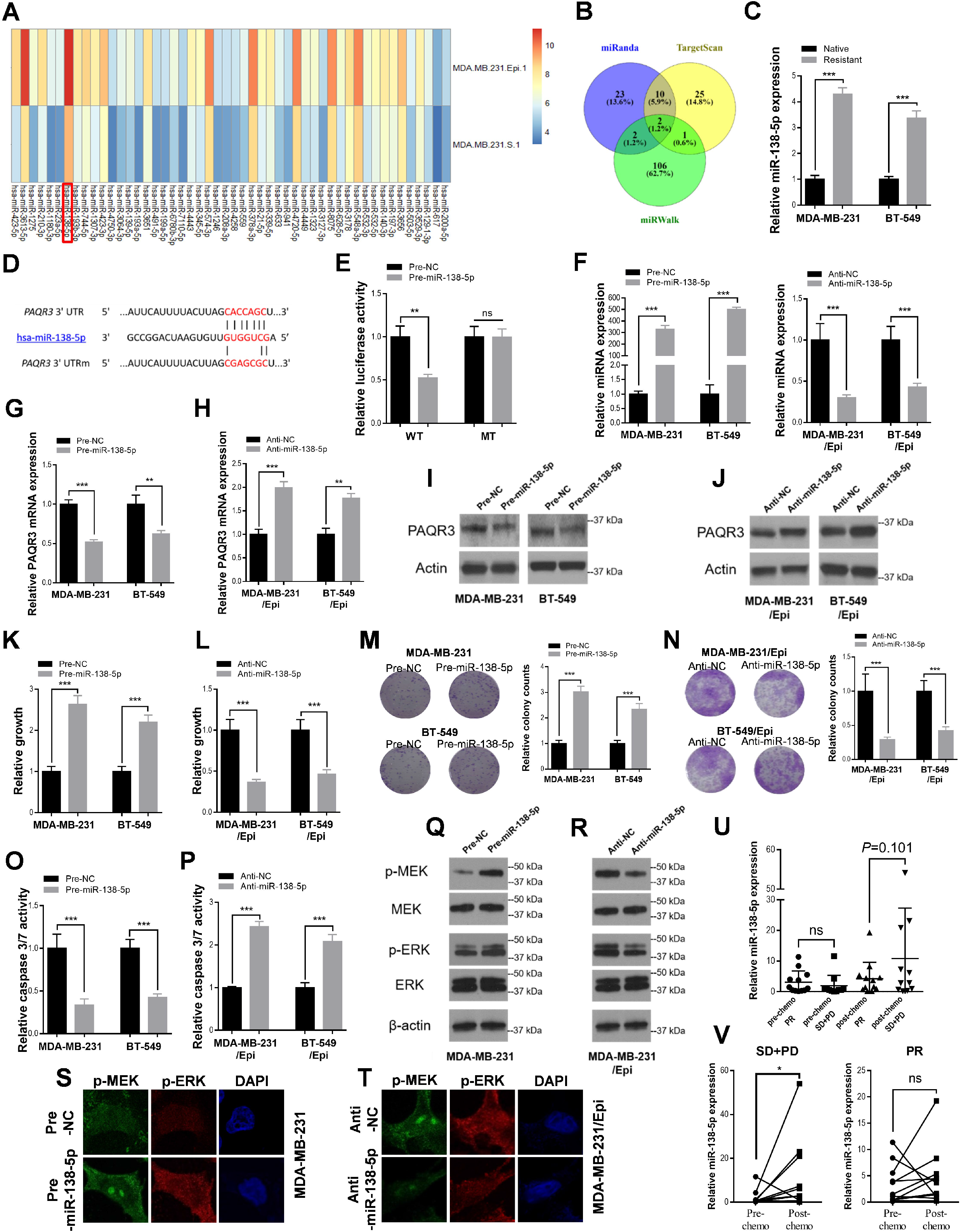
The upregulation of miR-138-5p was responsible for PAQR3 reduction-mediated resistance in epirubicin-resistant cells. (A) MiR-138-5p ranked top of the most upregulated miRNAs in MDA-MB-231/Epi cells, highlighted in red box. (B) MiR-138-5p was co-predicted targeting the 3’UTR of PAQR3 by miRanda, TargetScan and miRWalk databases. (C) The real-time PCR assay showed a 4.3- and 3.4-fold increase in miR-138-5p expression in MDA-MB-231/Epi and BT-549/Epi cells, when compared with native cells, respectively. (D) Illustration of the interaction between miR-138-5p and targeting sites within the 3’-UTR or mutant 3’-UTR of PAQR3. (E) Relative luciferase activity in MDA-MB-231 cells that were transfected with firefly luciferase reporters containing WT or mutant 3’-UTRs of PAQR3, pre-miR-138-5p, pre-NC. An average 47% decrease in luciferase activity was observed in WT reporter and pre-138-5p cotransfected cells, whereas no difference was detected in mutant plasmid transfected cells. (F) After artificially enforcing or suppressing miR-138-5p expression in native or resistant cells, the mRNA and protein levels of PAQR3 in native cells (MDA-MB-231 and BT-549) (G and I) and resistant cells (MDA-MB-231/Epi and BT-549/Epi) (H and J) were correspondingly changed. Pre-miR-138-5p caused epirubicin resistance in TNBC cells, characterized with 2.7- and 2.2-fold increase in cell viability after 72 hours epirubicin treatment (K), 3.0- and 2.4-fold increase in colony formation ability (M), 63% and 57% decrease in caspase 3/7 activity (O) for MDA-MB-231 and BT-549 cells, respectively. Inversely, miR-138-5p suppression by anti-miR-138-5p ameliorated resistance in MDA-MB-231/Epi and BT-549/Epi cells, with an average 63% and 54% decrease in cell viability (L), 71% and 58% decrease in colony numbers (N), and 2.4- and 2.1-fold increase in caspase3/7 activity (P). Pre-miR-138-5p enhanced phosphorylation of MEK1/2 and ERK1/2 as showed in western blot (Q) and IF (S) assays, whereas anti-miR-138-5p decreased the levels of phosphorylated MEK1/2 and ERK1/2 (R and T). (U) No alternation of miR-138-5p between partial response (PR), and stable or progressive disease (SD/PD) samples was achieved before chemotherapy (pre-chemo), whereas an increasing trend was observed in SD/PD samples after chemotherapy (post-chemo) when compared with PR tumors (2^-ΔΔCt^: 4.2 vs 10.8, *P*=0.101). (V) Levels of miR-138-5p in PR tumors did not change after chemotherapy, whereas miR-138-5p in tumors with SD/PD significantly elevated after chemotherapy (2^-ΔΔCt^: 1.9 vs 10.8, *P*=0.034). ns: no significance, * *P* < 0.05, ** *P* < 0.01, *** *P* < 0.001.

### MiR-138-5p suppression ameliorated epirubicin resistance in TNBC cells

MiR-138-5p has been reported associating with resistance in leukemia ^[8]^ and prostate cancer ^[9]^. In this study, we found pre-miR-138-5p was able to induce epirubicin resistance in TNBC cells, characterized with 2.7- and 2.2-fold increase in cell viability after 72 hours (Fig. 4K), 3.0- and 2.4-fold increase in colony formation ability (Fig. 4M), 63% and 57% decrease in caspase 3/7 activity (Fig. 4O) for MDA-MB-231 and BT-549 cells, respectively. Inversely, anti-miR-138-5p ameliorated resistance in resistant cells, with an average 63% and 54% decrease in cell viability (Fig. 4L), 71% and 58% decrease in colony counting (Fig. 4N), 2.4- and 2.1-fold increase in caspase3/7 activity for MDA-MB-231/Epi and BT-549/Epi cells (Fig. 4P). Pre-miR-138-5p enhanced phosphorylation of MEK1/2 and ERK1/2 in TNBC cells (Fig. 4Q and S), whereas anti-miR-138-5p decreased the levels of phosphorylated MEK1/2 and ERK1/2 in TNBC resistant cells (Fig. 4R and T).

The miR-138-5p expression in tumors with partial response (PR), and stable or progressive disease (SD/PD) was compared before and after chemotherapy. No difference between PR and SD/PD samples was found before chemotherapy, whereas an increasing trend was observed in SD/PD samples when compared with PR tumors after chemotherapy (2^-ΔΔCt^: 4.2 vs 10.8, *P* = 0.101) (Fig. 4U). In tumors with PR, miR-138-5p expression did not change after chemotherapy. Interestingly, miR-138-5p in SD/PD tumors significantly elevated after chemotherapy (2^-ΔΔCt^: 1.9 vs 10.8, *P* = 0.034) (Fig. 4V), which indicated a role of miR-138-5p in epirubicin resistance induction.

### The response to epirubicin-based chemotherapy could be predicted by PAQR3

Anthracyclines benefit for BC could be predicted with specific molecular markers. TOP2A, which acts as a target molecule of anthracyclines, is recognized relating with anthracyclines response ^[10, 11]^. Our previous finding indicated PAQR3 may predict the anthracyclines response in BC.

The baseline characteristics and pCR status of enrolled 114 ER negative patients were summarized in Table S3. A pCR rate of 14.0% was achieved, tumors with pCR were associated with distant metastasis-free survival (HR = 0.29, 95% CI = 0.10 to 0.84) (Fig. S1A). The associations of pCR status and demographics, clinical characteristics, transcriptional expression and gene amplification were described in Table 1. TOP2A amplification was expectedly associated with increased pCR rate. Interestingly, pCR rate was 21.0% and 7.0% in high- and low-PAQR3 tumors, the significant association of PAQR3 and pCR rate was observed (OR = 3.53; 95% CI, 1.06-11.72; *P* = 0.039), and maintained after adjusting other favorable predictors including TOP2A amplification (OR = 4.86, 95% CI = 1.13-20.87, *P* = 0.034). Moreover, PAQR3 was more closely associating with tumor response than TOP2A expression (OR = 1.00; 95% CI, 0.35-2.88; *P* = 1.000). The SPE, NPV, SEN and PPV of markers were summarized in Table S4. PAQR3 was more favorable than TOP2A expression in predicting tumor response. Moreover, NPV (0.93; 95% CI, 0.83-0.97) and SEN (0.75; 95% CI, 0.51-0.90) of PAQR3 were even higher than those based on TOP2A amplification (Fig. 5A).

**Figure 5.**
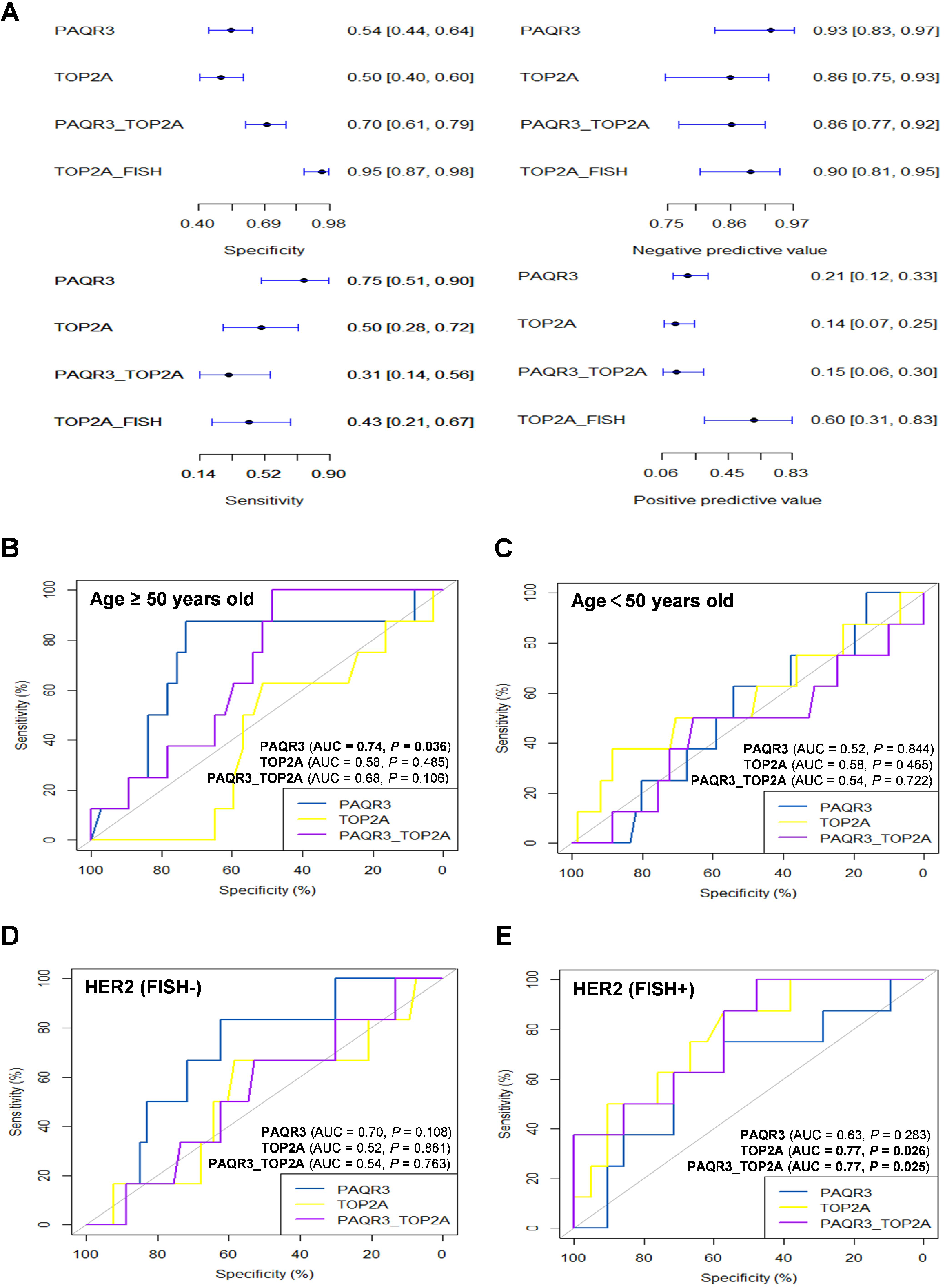
PAQR3 predicted tumor response to epirubicin. (A) The SPE, NPV, SEN and PPV were determined at the median for PAQR3 and TOP2A, and at the condition that tumors possessing both high levels of PAQR3 and TOP2A (PAQR3_TOP2A). The horizontal lines correspond to exact 95% CI. (B) In patients order than 50 years old, the AUCs were used to evaluate the prediction performance of PAQR3, TOP2A and PAQR3_TOP2A, PAQR3 possessed a better predictive value than TOP2A expression (0.74 vs 0.58), (C) whereas in patients younger than 50, predictive value by PAQR3 was comparable with TOP2A expression (D) In breast cancer without HER2 amplification, PAQR3 appeared better than TOP2A expression in epirubicin response prediction (0.70 vs 0.52), (E) no difference between PAQR3 and TOP2A was observed in HER2 amplified breast cancer.

**Table 1.** OR for pCR according to the demographical and clinical parameters, FISH and mRNA

| Characteristics | No. of Patients | Patients With pCR (%) | OR | 95% CI | P |
| --- | --- | --- | --- | --- | --- |
| **Univariate logistic regression** |  |  |  |  |  |
| **Age, years** |  |  | 1.65 | 0.57-4.77 | 0.356 |
| ≤ 50 (Ref) | 69 | 8 (11.6) |  |  |  |
| > 50 | 45 | 8 (17.8) |  |  |  |
| **Tumor size** |  |  | 1.18 | 0.30-4.63 | 0.810 |
| T1-T2 (Ref) | 95 | 13 (13.7) |  |  |  |
| T3-T4 | 19 | 3 (15.8) |  |  |  |
| **Nodal status** |  |  | 1.47 | 0.50-4.37 | 0.484 |
| N0 (Ref) | 52 | 6 (11.5) |  |  |  |
| N1-3 | 62 | 10 (16.1) |  |  |  |
| **Histologic grade** |  |  | 1.01 | 0.26-3.96 | 0.985 |
| G1-G2 (Ref) | 22 | 3 (13.6) |  |  |  |
| G3 | 87 | 12 (13.8) |  |  |  |
| **HER2_FISH** |  |  | 3.37 | 1.04-10.87 | **0.043** |
| No amplification (Ref) | 59 | 6 (10.2) |  |  |  |
| Amplification | 29 | 8 (27.6) |  |  |  |
| **TOP2A_FISH** |  |  |  |  |  |
| No amplification (Ref) | 77 | 8 (10.4) | 12.94 | 3.00-55.80 | **<0.001** |
| Amplification | 10 | 6 (60.0) |  |  |  |
| Deletion (Ref) | 10 | 2 (20.0) | 0.39 | 0.07-2.29 | 0.299 |
| Normal | 67 | 6 (9.0) |  |  |  |
| Normal (Ref) | 67 | 6 (9.0) | 15.25 | 3.34-69.58 | **<0.001** |
| Amplification | 10 | 6 (60.0) |  |  |  |
| **TOP2A_mRNA** |  |  | 1.00 | 0.35-2.88 | 1.000 |
| Low (Ref) | 57 | 8 (14.0) |  |  |  |
| High | 57 | 8 (14.0) |  |  |  |
| **PAQR3_mRNA** |  |  | 3.53 | 1.06-11.72 | 0.039 |
| Low (Ref) | 57 | 4 (7.0) |  |  |  |
| High | 57 | 12 (21.0) |  |  |  |
| **Multivariate logistic regression** |  |  |  |  |  |
| **TOP2_mRNA** |  |  | 0.56 | 0.12-2.76 | 0.479 |
| **PAQR3_mRNA** |  |  | 4.86 | 1.13-20.87 | **0.034** |
| **HER2_FISH** |  |  | 0.90 | 0.14-5.49 | 0.897 |
| **TOP2A_FISH** |  |  | 24.02 | 1.96-294.00 | **0.013** |

TOP2A and PAQR3 mRNA binary values were determined at the median of whole cohort. OR, odds ratio; CI, confidence interval; pCR, pathologic complete response; FISH, fluorescent in situ hybridization; TOP2A, topoisomerase II-α; PAQR3, pogestin and adipoQ receptor family member 3; HER2, human epidermal growth factor receptor 2.

### PAQR3 better predicted tumor response to epirubicin in old patients

BC in elderly may not benefit anthracyclines-based chemotherapy, which may cause severe and irreversible cardiotoxicity. Therefore, there has been a growing interest to identify low-risk patients who may be candidates for de-intensification of anthracyclines-based chemotherapy without compromising survival.

For patients older than 50 years old, PAQR3 exhibited an improved predictive value for tumor response to epirubicin, when compared with its performance in the whole cohort. The achieved pCR rate in high-PAQR3 group increased from 21.0% to 35.0%, was much higher than that in low-PAQR3 tumors (4.5%). OR increased from 3.53 to 12.92 (95% CI, 1.43-116.78, *P* = 0.023). The SPE, NPV, SEN and PPV increased from 0.54 to 0.65 (95% CI, 0.49-0.78), 0.93 to 0.96 (95% CI, 0.80-0.99), 0.75 to 0.88 (95% CI, 0.53-0.98), and 0.21 to 0.35 (95% CI, 0.18-0.57), respectively (Table 2). Moreover, PAQR3 expression was lower in patients older than 70 as compared to others (50-70 years) (Fig. S1B), suggesting some patients (>70 years) may be qualified for de-intensification of epirubicin. The AUC assessing the prediction performance of PAQR3 was higher than that of TOP2A (0.74 vs 0.58) (Fig. 5B). However, in patients younger than 50, we did not observe an increased predictive value for PAQR3 (Fig. 5C) (Table 2).

**Table 2.**
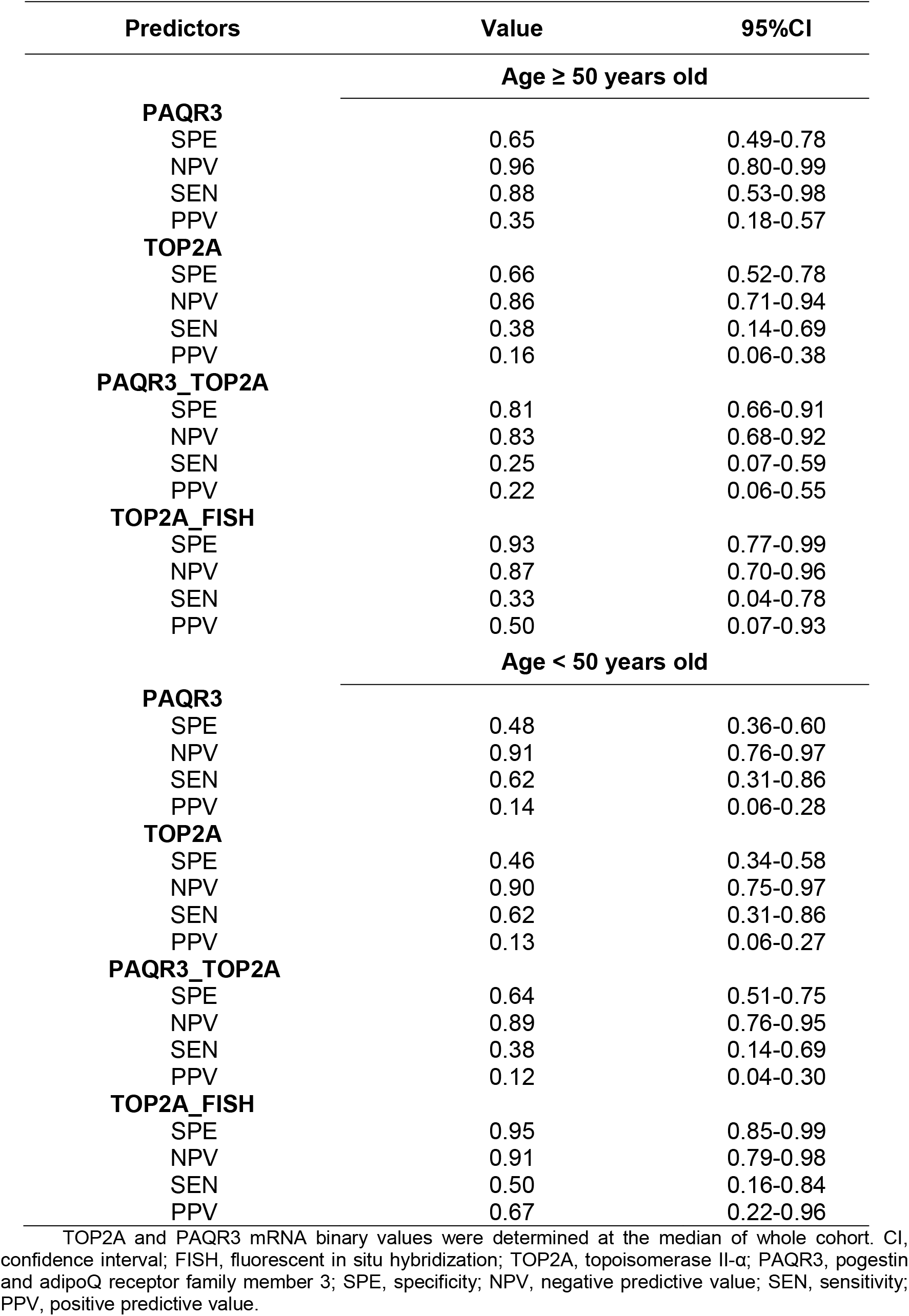
SPE, NPV, SEN and PPV stratified by age

Human epithelial receptor 2 (HER2) also functions as a vital degerminator in BC growth, progression and response to treatment. In BC without HER2 amplification, PAQR3 appeared having the superiority over TOP2A expression in epirubicin response prediction (0.70 vs 0.52) (Fig. 5D), whereas TOP2A expression was better in HER2-amplified BC (Fig. 5E).

### The PAQR3_TOP2A predicted tumor response after anthracyclines- or taxane/anthracyclines-based chemotherapy

We also sought to see if PAQR3_TOP2A could better predict the response to epirubicin. Subsequently, PAQR3_TOP2A possessed higher SPE (0.70; 95% CI, 0.61-0.79), as compared to either PAQR3 or TOP2A alone. Also, PAQR3_TOP2A had a high NPV (0.86, 95% CI: 0.77-0.92) (Fig. 5A), which enables identifying patients qualified for de-intensification of epirubicin.

We further detected if PAQR3_TOP2A was favorable in predicting the response to anthracyclines- or taxane/anthracyclines-based chemotherapy in ER negative patients. The SPE, NPV, SEN, PPV and OR calculated with PAQR3, TOP2A and PAQR3_TOP2A were summarized in Table S5. Similarly, PAQR3_TOP2A induced an elevated SPE (ranging from 0.74 to 0.79) across all datasets (Fig. S2A), when compared with either PAQR3 or TOP2A alone (Table S5). Also, PAQR3_TOP2A had a high NPV ranging from 0.54 to 0.78 (Fig. S2B). PAQR3_TOP2A was significantly associated with pCR rate in patients receiving anthracyclines-based chemotherapy, the ORs were 4.04 (95% CI: 1.22-13.39) and 3.89 (95% CI: 1.18-12.85), whereas no statistical associations were observed in patients receiving taxane/anthracyclines-based chemotherapy (Fig. S2C).

ER positive BC tends to be less responsive to anthracyclines-based chemotherapy, therefore, it is plausible to identify patients qualified for de-intensification of anthracyclines-based chemotherapy in ER positive patients (Table S6). Similarly, PAQR3_TOP2A induced an elevated SPE, ranging from 0.68 to 0.77 (Fig. S2D), as compared to either PAQR3 or TOP2A alone. Due to the favorable NPV from both PAQR3 and TOP2A, PAQR3_TOP2A consistently generated a high NPV, ranging from 0.79 to 0.92 (Fig. S2E). PAQR3_TOP2A was only significantly associated with pCR rate in patients receiving taxane/anthracycline-based chemotherapy (OR = 2.97, 95% CI: 1.37-6.45) (Fig. S2F).

## Discussion

Normally, MAPK signaling cascade is carefully orchestrated, but in cancer, the aberrant activation is always initiated by mutation, which led to an 80-fold increase in doxorubicin IC_50_ ^[12]^. Different from prostate cancer and melanoma harboring a high average frequency of KRas or V600E BRaf mutation, BC rarely has MAPK pathway components mutations ^[13]^, this in part leads to the less extensive study on MAPK pathway targeting therapy in BC. One clinical trial combatively using MEK1/2 inhibitor and fulvestrant failed to improve patients’ outcomes in BC progressing after aromatase inhibitor therapy ^[14]^, implying us MEK1/2 inhibitor may only produce a promising result in tumors with MAPK pathway aberrant activation. Interestingly, MAPK pathway in BC could be activated by anthracyclines ^[3]^, in this study, we further observed MAPK pathway activation in epirubicin-resistant BC, abrogation with either MEK1/2 or ERK1/2 inhibitor ameliorated epirubicin resistance.

PAQR3 reduction was identified as the main cause for MAPK activation in epirubicin-resistant TNBC cells, as PAQR3 abates ERK1/2 phosphorylation via trapping the upstream phosphokinase Raf to Golgi ^[5]^. We found PAQR3 possessed a role of ameliorating epirubicin resistance, besides its inhibitory role in malignancies growth and progression ^[6, 15]^. PAQR3 could be regulated by miRNAs induced epigenetic silencing ^[16]^, miR-138-5p, one of the most upregulated miRNAs in MDA-MB-231/Epi cells, was experimentally proved targeting PAQR3. MiR-138-5p was previously reported inducing resistance to docetaxel in prostate cancer ^[17]^ and gefitinib in NSCLC ^[18]^, it was also involved in BC susceptibility ^[19]^ and tumor progression ^[20]^. Our finding in current study facilitates a more comprehensive understanding miR-138 in TNBC.

Identifying BC patients not sensitive to anthracyclines-based chemotherapy is so meaningful. Generally, chemoresistance could be determined through evaluating tumor response after neo-adjuvant chemotherapy, however, for patients receiving breast mastectomy and adjuvant chemotherapy, tumor response cannot be easily predicted. Therefore, exploring a more applicable marker is important for chemoresistance prediction. Here, we first identified PAQR3 possessed a predictive value in determining the pCR rate achieved by anthracyclines-based chemotherapy in BC. TOP2A is the target molecule of anthracyclines, amplification or expression of TOP2A have been considered predictor for anthracyclines-based chemotherapy. The Collaborative Study Group of Scientific Research of the Japanese Breast Cancer Society proved TOP2A positive BC had a significant higher pCR rate as compared to negative cancer after epirubicin treatment ^[21]^. Excitingly, PAQR3 was more favorable than TOP2A expression in predicting pCR status, the NPV and SEN of PAQR3 were even higher than TOP2A amplification, the most reliable marker for predicting the response to anthracyclines-based chemotherapy.

Moreover, the better predictive value of PAQR3 in old patients suggests it can be used to identify patients more likely benefit from epirubicin containing regimens, also helps non-response old patients avoid epirubicin-caused cardiotoxicity. Interestingly, patients older than 70 years had the lower PAQR3 expression suggested they may be qualified for de-intensification of epirubicin. In patients with HER2 non-amplified BC, PAQR3 had a superiority over TOP2A expression in predicting the epirubicin response, whereas TOP2A expression was better in HER2 amplified BC, the co-amplification of TOP2A in 40% of HER2 amplified BC may explain this observed disparity ^[22]^.

PAQR3_TOP2A consistently increased SPE in both ER positive and negative patients, which promotes precisely identifying patients not benefit from anthracyclines. NPV represents the accuracy of negative results produced by the screening test, in this study, it was the probability that patients with no-pCR prediction test result indeed did not have pCR after anthracyclines-based chemotherapy. The favorable NPV of PAQR3_TOP2A in both ER positive and negative patients proved the reliability of using PAQR3_TOP2A for chemotherapeutic response prediction, through avoiding false negatives.

Collectively, with the less biased methodologies, we discovered miR-138-5p induced PAQR3 reduction causes epirubicin resistance in TNBC cells via MAPK cascades activation. PAQR3 is a potential predictor for tumor response to anthracyclines in clinic, but still needs a large-scale sample to confirm, especially at the protein level.

## Materials and Methods

### Chemicals and antibodies

PD98059 (#9900, Cell Signaling Technology, MA, USA) used at 100 μM, SCH772984 (#S7101, Selleckchem, TX, USA) used at 10 μM. p-ERK1/2 (#5726, Mouse, western blot (WB) 1:1000, immunofluorescence (IF) 1:200), p-MEK1/2 (#2338, Rabbit, WB 1:2000), p-MEK1 (#9127, Rabbit, IF 1:100), ERK1/2 (#9102, Rabbit, WB 1:1000) and MEK1/2 (#4694, Mouse, WB 1:1000) antibodies were purchased from Cell Signaling Technology, MA, USA. Mouse β-actin antibody (#SC517582, Santa Cruz, TX, USA) was used at 1:1000, Rabbit PAQR3 antibody (#ab174327, Abcam, Cambridge, UK) was used at 1:200.

### Cell culture and epirubicin resistance induction

Human breast cancer cell lines (MDA-MB-231, MCF-7, SK-BR-3 and BT-549), and HEK293T cells were purchased from American Type Culture Collection (Manassas, VA, USA). Epirubicin resistant cell lines were generated as previously described by Braunstein et al. ^[23]^, briefly, cells were stepwise exposed to an increasing concentrations of epirubicin (Pfizer Pharmaceuticals, Wuxi, China) for 12 months, with an initial concentration set at 5 nM until 100 nM, each increasing dose was given when the cells were in proliferation without significant death. Resistant cells were maintained in culture with 100 nM epirubicin, which was withdrawn from culture medium for at least 2 passages before processing an experiment All cells were maintained in RPMI 1640 medium (Gibco, MD, USA) supplemented with 10% fetal bovine serum (Invitrogen, CA, USA), 100 U/mL penicillin, and 100 μg/mL streptomycin, at 37°C in a humidified incubator with 5% CO_2_ injection. Mycoplasma testing was carried out routinely every 6-8 weeks.

### Cell viability and colony formation measurement

Cell viability was measured with Cell Counting Kit-8 (CCK-8) (Dojindo, Tokyo, Japan). In brief, 3×10^3^ cells in quintuplicate were seeded into 96-well plate for each group, CCK-8 was added and incubated with cells for one hour according to manufacturer’s instructions at indicated time points, followed by measurement with a microplate reader at 450 nm wavelength. The half maximal inhibitory concentration was calculated after an increasing concentration of epirubicin treatment for 72 hours. For colony formation assay, 3×10^3^ cells were seeded in six-well plates and maintained in complete medium for two weeks, the surviving colonies were fixed with methanol, stained with crystal violet, and then counted. 0.5 μM and 0.3 μM epirubicin were utilized in cell proliferation and colony formation assays, respectively, when epirubicin resistance of tumor cells was evaluated. Each experiment was repeated three times, independently.

### Plasmid construction and miRNA transfection

A complementary DNA expression vector encoding full length of PAQR3 was cloned into pcDNA3.1(+) plasmid as previously reported ^[24]^, the resultant plasmid was then sequenced. Lipofectamine 2000 (Invitrogen, CA, USA) was utilized to mediate plasmid transfection. Clones with stable expression were selected by G418 (Thermo Fisher Scientific, MA, USA). The transfected cells were selected with 800 μg/mL G418, and maintained in culture medium containing 600 μg/mL after the control cells died. The RKTG short hairpin RNA (shRNA) construct was generated using a lentiviral system. In short, an annealed small interfering RNA cassette with a targeting sequence of GGACAACCCGUACAUCACC for RKTG and scramble sequences were inserted into the pLKO.1.-puro vector (Addgene, MA, USA) downstream of the U6 promoter Lentiviruses were obtained by co-transfecting a mixture of the indicated shRNA plasmid, the envelope plasmid (pMD2.G) and the packaging plasmid (pCMV-deltaR8.91) (System Biosciences, CA, USA) into HEK293T cells according to the manufacturer’s instructions. Synthetic pre-miR-138-5p, anti-miR-138-5p and scrambled negative control RNA (GenePharma, Shanghai, China) were transfected following manufacturer’s instructions.

### RNA extraction and real-time PCR

Total RNA from tumor cells was isolated using TRIzol^®^ reagent (Invitrogen, CA, USA) as instructed by the provided protocol. The mRNA and miRNA reverse transcription were performed within a 5×All-In-One RT MasterMix kit (Applied Biological Materials, BC, Canada) and TaqMan microRNA Reverse Transcription Kit (Thermo Fisher, MD, USA). Real-time PCR was performed using SYBR^®^ Green Master Mix (Applied Biosystems, CA, USA) according to the manufacturer’s protocol and reacted in 7500 real-Time PCR system. β-actin and U6 were used as the references for PAQR3 and miR-138-5p, respectively. The primers were listed in Table S1, each real-time PCR experiment was independently repeated at least three times.

### Protein extraction and immunoblotting

The whole cell lysate was extracted by RIPA lysis and extraction buffer (Thermo Fisher Scientific, CA, USA) with protease inhibitor cocktail (Sigma-Aldrich, MO, USA) and phosSTOP^TM^ (Sigma-Aldrich, MO, USA). BCA protein assay kit (Thermo Fisher Scientific, MA, USA) was used measuring protein concentration per manufacturer instructions. 30 μg of total protein for each sample was separated via sodium dodecyl sulphate/polyacrylamide gel electrophoresis (SDS-PAGE) and transferred onto 0.45 μm pore-size nitrocellulose membrane (Life Science, CA, USA). Membrane was then blocked with 5% non-fat milk in tris-buffered saline with 0.5% Tween at room temperature for one hour, followed by overnight incubation with primary antibody at 4 °C and an hour incubation with secondary antibody at room temperature. The enhanced chemiluminescence (Thermo Fisher Scientific, CA, USA) was used to produce light emission.

### Immunofluorescence (IF)

Cells were fixed with 4 % paraformaldehyde for 15 minutes, then blocked with the buffer containing 5% goat serum (Abcam, MA, USA) and 0.3% Triton™ X-100 (Sigma-Aldrich, MO, USA) for one hour at room temperature. After overnight primary antibody incubation at 4 °C, cells were incubated with fluorochrome-conjugated secondary antibody (Thermo Fisher Scientific, CA, USA) in dark for one hour, followed by 1 μg/ml DAPI (Thermo Fisher Scientific, CA, USA) staining for 10 minutes at room temperature and Prolong^®^ Gold Antifade Reagent mounting (Thermo Fisher Scientific, CA, USA).

### Caspase 3/7 activity assay

Breast cancer cells were plated in quadruplicate in 96-well plates. After a 24-hour incubation, the cells were then treated with 0.5 mg/ml epirubicin for 72 hours. Caspase 3/7 assay reagents (Promega, WI, USA) were added to each well according to the manufacturer’s instructions (ratio of 1:4), immunofluorescence was measured, the excitation and emission wavelengths were set as 485 and 530 nm, respectively.

Differential expression genes (DEGs) identification and Gene set enrichment analysis (GSEA)

The transcriptional expression data of epirubicin-resistant and native breast cancer cells were retrieved from GSE54326 dataset and processed with R software (version: 3.6.1, https://www.r-project.org/), the data of this illumina microarray was normalized with a method called “quantile” under the package “lumi”. DEGs between resistant and native cells were identified with linear model under the package “limma”. The empirical Bayes method was used to borrow information between genes. Genes in negative regulation of MAPK pathway were indexed in AmiGO2 (http://geneontology.org/) and summarized in Table S2. The volcano plot was used to present differential expression of these genes in MDA-MB-231, MCF-7 and SK-BR-3. GSEA was based on GSE54326 and processed with GSEA software (http://software.broadinstitute.org/gsea/index.jsp) ^[25]^.

### Luciferase activity assay

The 3’UTR of human PAQR3 was cloned into pmirGLO vector (Promega, Madison, WI, USA), and then validated by sequencing. The 3’UTR sequence which interacted with seed sequence of miR-138-5p was mutated to further test the biding specificity of miR-138-5p on PAQR3. MDA-MB-231 cells were co-transfected with luciferase reporter plasmid and pre-miR-138-5p/pre-NC. The Renilla luciferase plasmid (Promega, Madison, WI, USA) was used as transfection control. The luciferase activity was measured with Dual-Luciferase Reporter Assay System (Promega, Madison, WI, USA), after harvesting the 48 hours transfected cells.

### In vivo breast cancer studies

All in vivo studies were performed with xenograft tumor model. Native and epirubicin resistant tumor cells (3×10^6^ cells) were injected subcutaneously into the dorsal right flank of 5-week-old female athymic mice, when the tumor volume reached at approximate 100 mm^3^, PD98059 (10 mg/kg) or epirubicin (10 mg/kg) was administered to the mice in a 0.2 ml intraperitoneal injection with a 3-day interval for three weeks. The tumor volumes were measured every two to three days, tumor volume was calculated according to the formula: volume = 0.5 × width^2^ × length. The tumors were weighed after removal. The experimental procedures were approved by the Animal Ethics Committee of The First Affiliated Hospital of Chongqing Medical University. All the mice were housed according to the national and institutional guidelines for humane animal care.

### Study population

The clinical and transcriptional data of 114 ER negative BC patients were available and retrieved from the prospective multicentric TOP trial ^[26]^ (http://www.ncbi.nlm.nih.gov/geo/). As previously described, patients with locally advanced and inflammatory disease received epirubicin monotherapy (100 mg/m^2^) as the neoadjuvant chemotherapy, with four cycles every three weeks for patients with early BC, and a dose-dense schedule of six cycles every two weeks for patients with locally advanced and inflammatory disease. After chemotherapy complement, tumor response to epirubicin was evaluated, pathological complete response (pCR) was defined as no residual invasive cancerous tissue, or existence of in situ carcinoma without invasion in breast and axillary nodes.

Other cohorts receiving anthracyclines- or taxane/anthracyclines-based chemotherapy were used to evaluate the combination of PAQR3 and TOP2A (PAQR3_TOP2A) in predicting the tumor response ^[27–31]^ (http://www.ncbi.nlm.nih.gov/geo/).

### Statistical analysis

Kaplan-Meier method was used for estimating the survival rates, the difference was analyzed by log-rank test, hazard ratio (HR) was used to compare the probability of events between groups. The sensitivity (SEN), specificity (SPE), positive predictive value (PPV) and negative predictive value (NPV) were determined at the threshold of gene medium expression The area under the curve (AUC) was applied to assess the prediction performance of PAQR3, TOP2A and PAQR3_TOP2A. AUC was estimated with the concordance index, the confidence interval (CI) and significance being estimated assuming asymptotic normality. Odds ratio (OR) was calculated to compare pCR rates between groups defined in this study. Logistic regression was used for simultaneously assessing multivariable factors predicting the pCR rates. The Student’s t-test or paired t-test was performed to evaluate the differences between two groups. We used the median value of gene transcriptional expression to group the patients. Frequency counts were compared between groups using the two-tailed Fisher’s exact test. The statistical analysis was performed with GraphPad Prism 5.0 (GraphPad Software, CA, USA) and R version 3.6.1 (https://www.r-project.org/).

## Acknowledgements

This work was supported by the National Natural Science Foundation of China (31420103915 to G. R).

## Author contributions

J.H. and G.R. conceived the project, G.R. supervised the experiments. J.H., S.Z., Y.X., X.W. and T.X. performed the experiments. J.H. and S.Z. performed bioinformatics analysis. J.H., H.L. and L.K. performed statistical analysis. J.H. and Y.X. wrote the manuscript with help form all the authors. All authors reviewed the manuscript.

---

**Figure S1. pCR was associated with favorable survival.** (A) Tumors with pCR were associated with improved distant metastasis-free survival (HR = 0.29, 95% CI = 0.10 to 0.84, *P* = 0.023), and appeared having a favorable overall survival (HR = 0.28, 95% CI = 0.07 to 1.072, *P* = 0.063). (B) PAQR3 expression was lower in patients older than 70, as compared to other patients older than 50, but less than 70 years old. *** *P* < 0.001.

**Figure S2. PAQR3 predicted tumor response to anthracyclines- or taxane/anthracyclines-based chemotherapy.** In ER negative breast cancer, PAQR3_TOP2A had an elevated SPE ranging from 0.74 to 0.79 (A), a high NPV ranging from 0.54 to 0.78 (B), and was significantly associated with pCR rate in patients receiving anthracyclines-based chemotherapy (ORs = 4.04 (95% CI: 1.22-13.39) and 3.89 (95% CI: 1.18-12.85)) across all cohorts (C). In ER positive breast cancer, PAQR3_TOP2A induced an elevated SPE ranging from 0.68 to 0.77 (D), consistently generated a high NPV ranging from 0.79 to 0.92 (E), and was only significantly associated with pCR rate in patients receiving taxane/anthracycline-based chemotherapy (OR = 2.97, 95% CI: 1.37-6.45) (F).

