## Supplemental Table 5 for "MAPK Pathway Inhibition as A Rational Therapeutic Strategy for MiR-138-5p/PAQR3 Dysregulation-mediated Epirubicin Resistance in Triple-negative Breast Cancer"

### Table S5. SPE, NPV, SEN, PPV and OR in ER negative breast cancer

| Predictors | Value | 95%CI | Value | 95%CI |
|---|---|---|---|---|
| | Wessel et al. Anthracyclines-based (n=57 (33 pCRs)) | | Iwanoto et al. Anthracyclines-based (n=55 (18 pCRs)) | |
| **SPE** | | | | |
| PAQR3 | 0.67 | 0.45-0.84 | 0.51 | 0.34-0.68 |
| TOP2A | 0.62 | 0.41-0.81 | 0.62 | 0.45-0.78 |
| PAQR3_TOP2A | 0.79 | 0.60-0.91 | 0.76 | 0.60-0.87 |
| **NPV** | | | | |
| PAQR3 | 0.57 | 0.37-0.75 | 0.70 | 0.50-0.86 |
| TOP2A | 0.54 | 0.34-0.72 | 0.85 | 0.66-0.96 |
| PAQR3_TOP2A | 0.54 | 0.38-0.70 | 0.78 | 0.62-0.88 |
| **SEN** | | | | |
| PAQR3 | 0.64 | 0.45-0.80 | 0.55 | 0.31-0.78 |
| TOP2A | 0.61 | 0.42-0.77 | 0.78 | 0.52-0.94 |
| PAQR3_TOP2A | 0.52 | 0.35-0.67 | 0.56 | 0.34-0.75 |
| **PPV** | | | | |
| PAQR3 | 0.72 | 0.53-0.87 | 0.36 | 0.19-0.56 |
| TOP2A | 0.69 | 0.49-0.85 | 0.50 | 0.31-0.69 |
| PAQR3_TOP2A | 0.77 | 0.57-0.90 | 0.53 | 0.32-0.73 |
| **OR** | | | | |
| PAQR3 | 3.50 | 1.16-10.58 | 1.32 | 0.43-4.09 |
| TOP2A | 2.56 | 0.87-7.56 | 5.75 | 1.57-20.99 |
| PAQR3_TOP2A | 4.04 | 1.22-13.39 | 3.89 | 1.18-12.85 |
| | Booser et al. Taxane/Anthracyclines-based (n=219 (73 pCRs)) | | Prat et al. Taxane/Anthracyclines-based (n=56 (15 pCRs)) | |
| **SPE** | | | | |
| PAQR3 | 0.48 | 0.40-0.56 | 0.54 | 0.37-0.69 |
| TOP2A | 0.53 | 0.45-0.62 | 0.51 | 0.35-0.67 |
| PAQR3_TOP2A | 0.74 | 0.66-0.80 | 0.76 | 0.61-0.86 |
| **NPV** | | | | |
| PAQR3 | 0.64 | 0.54-0.73 | 0.81 | 0.62-0.94 |
| TOP2A | 0.72 | 0.63-0.80 | 0.75 | 0.55-0.89 |
| PAQR3_TOP2A | 0.68 | 0.60-0.75 | 0.76 | 0.61-0.86 |
| **SEN** | | | | |
| PAQR3 | 0.47 | 0.35-0.59 | 0.67 | 0.38-0.88 |
| TOP2A | 0.59 | 0.47-0.70 | 0.53 | 0.27-0.79 |
| PAQR3_TOP2A | 0.30 | 0.21-0.41 | 0.33 | 0.15-0.58 |
| **PPV** | | | | |
| PAQR3 | 0.31 | 0.22-0.40 | 0.34 | 0.18-0.54 |
| TOP2A | 0.39 | 0.30-0.48 | 0.29 | 0.13-0.49 |
| PAQR3_TOP2A | 0.37 | 0.26-0.49 | 0.33 | 0.15-0.58 |
| **OR** | | | | |
| PAQR3 | 0.80 | 0.46-1.41 | 2.32 | 0.67-7.97 |
| TOP2A | 1.64 | 0.93-2.90 | 1.20 | 0.37-3.92 |
| PAQR3_TOP2A | 1.23 | 0.66-2.28 | 1.55 | 0.43-5.62 |

TOP2A and PAQR3 mRNA binary values were determined at the median. PAQR3_TOP2A was defined as enrichment with both high TOP2A and PAQR3 mRNA. CI, confidence interval; TOP2A, topoisomerase II-α; PAQR3, pogestin and adipoQ receptor family member 3; SPE, specificity; NPV, negative predictive value; SEN, sensitivity; PPV, positive predictive value; OR, odds ratio.
