## Supplemental Table 6 for "MAPK Pathway Inhibition as A Rational Therapeutic Strategy for MiR-138-5p/PAQR3 Dysregulation-mediated Epirubicin Resistance in Triple-negative Breast Cancer"

### Table S6. SPE, NPV, SEN, PPV and OR in ER positive breast cancer

| Predictors | Value | 95%CI | Value | 95%CI |
|---|---|---|---|---|
| | Wessel et al. Anthracyclines-based (n=118 (24 pCRs)) | | Iwanoto et al. Anthracyclines-based (n=41 (7 pCRs)) | |
| **SPE** | | | | |
| PAQR3 | 0.50 | 0.39-0.60 | 0.44 | 0.27-0.62 |
| TOP2A | 0.52 | 0.41-0.62 | 0.50 | 0.32-0.68 |
| PAQR3_TOP2A | 0.69 | 0.59-0.78 | 0.68 | 0.51-0.81 |
| **NPV** | | | | |
| PAQR3 | 0.80 | 0.67-0.89 | 0.75 | 0.51-0.91 |
| TOP2A | 0.83 | 0.71-0.92 | 0.89 | 0.67-0.99 |
| PAQR3_TOP2A | 0.81 | 0.71-0.88 | 0.79 | 0.62-0.90 |
| **SEN** | | | | |
| PAQR3 | 0.50 | 0.29-0.71 | 0.28 | 0.03-0.71 |
| TOP2A | 0.58 | 0.37-0.78 | 0.71 | 0.29-0.96 |
| PAQR3_TOP2A | 0.38 | 0.21-0.57 | 0.14 | 0.03-0.51 |
| **PPV** | | | | |
| PAQR3 | 0.20 | 0.11-0.33 | 0.09 | 0.01-0.30 |
| TOP2A | 0.24 | 0.14-0.37 | 0.23 | 0.08-0.45 |
| PAQR3_TOP2A | 0.24 | 0.13-0.39 | 0.08 | 0.01-0.35 |
| **OR** | | | | |
| PAQR3 | 1.00 | 0.41-2.45 | 0.32 | 0.05-1.86 |
| TOP2A | 1.52 | 0.61-3.77 | 2.50 | 0.42-14.71 |
| PAQR3_TOP2A | 1.34 | 0.53-3.43 | 0.35 | 0.04-3.26 |
| | Booser et al. Taxane/Anthracyclines-based (n=285 (30 pCRs)) | | Prat et al. Taxane/Anthracyclines-based (n=37 (7 pCRs)) | |
| **SPE** | | | | |
| PAQR3 | 0.50 | 0.44-0.56 | 0.50 | 0.31-0.69 |
| TOP2A | 0.53 | 0.47-0.60 | 0.57 | 0.37-0.74 |
| PAQR3_TOP2A | 0.77 | 0.72-0.82 | 0.77 | 0.59-0.88 |
| **NPV** | | | | |
| PAQR3 | 0.90 | 0.84-0.94 | 0.83 | 0.59-0.96 |
| TOP2A | 0.96 | 0.91-0.98 | 0.94 | 0.73-1.00 |
| PAQR3_TOP2A | 0.92 | 0.88-0.95 | 0.85 | 0.68-0.94 |
| **SEN** | | | | |
| PAQR3 | 0.53 | 0.34-0.72 | 0.57 | 0.18-0.90 |
| TOP2A | 0.80 | 0.61-0.92 | 0.86 | 0.42-1.00 |
| PAQR3_TOP2A | 0.47 | 0.30-0.64 | 0.43 | 0.16-0.75 |
| **PPV** | | | | |
| PAQR3 | 0.11 | 0.06-0.17 | 0.21 | 0.06-0.46 |
| TOP2A | 0.17 | 0.11-0.24 | 0.32 | 0.13-0.56 |
| PAQR3_TOP2A | 0.19 | 0.12-0.30 | 0.30 | 0.11-0.60 |
| **OR** | | | | |
| PAQR3 | 1.15 | 0.54-2.46 | 1.33 | 0.25-7.01 |
| TOP2A | 4.57 | 1.81-11.56 | 7.85 | 0.84-73.46 |
| PAQR3_TOP2A | 2.97 | 1.37-6.45 | 2.46 | 0.44-13.75 |

TOP2A and PAQR3 mRNA binary values were determined at the median. PAQR3_TOP2A was defined as enrichment with both high TOP2A and PAQR3 mRNA. CI, confidence interval; TOP2A, topoisomerase II-α; PAQR3, pogestin and adipoQ receptor family member 3; SPE, specificity; NPV, negative predictive value; SEN, sensitivity; PPV, positive predictive value; OR, odds ratio.
