## Supplemental Table 1 for "MAPK Pathway Inhibition as A Rational Therapeutic Strategy for MiR-138-5p/PAQR3 Dysregulation-mediated Epirubicin Resistance in Triple-negative Breast Cancer"

**Table S1** Oligonucleotide primers for real-time PCR

| Gene | Primers |
|---|---|
| PAQR3 | Forward: CAAGTGCGTCCAGAGAAGATT |
| | Reverse: TCTGACCGATGGCAGGAAAAA |
| PPP5C | Forward: AAGACTCAGGCCAATGACTACT |
| | Reverse: CGCGTAGCCATAGCACTCAG |
| PPP2R4 | Forward: CCGGGTGGATGACCAAATAGC |
| | Reverse: GAACTGCCCCAGATGAAGGG |
| DUSP1 | Forward: ACCACCACCGTGTTCAACTTC |
| | Reverse: TGGGAGAGGTCGTAATGGGG |
| DUSP2 | Forward: TGCCCCAACCACTTTGAGG |
| | Reverse: AGTCAATGAAGCCTATGGCCT |
| DUSP4 | Forward: GGCATCACGGCTCTGTTGAAT |
| | Reverse: GTCGGCCTTGTGGTTATCTTC |
| DUSP5 | Forward: GCCAGCTTATGACCAGGGTG |
| | Reverse: GTCCGTCGGGAGACATTCAG |
| DUSP6 | Forward: GAAATGGCGATCAGCAAGACG |
| | Reverse: CGACGACTCGTATAGCTCCTG |
| DUSP8 | Forward: GTCCCCATCAACGACAACTAC |
| | Reverse: CAGTGGACGATGACTTGGCAG |
| DUSP9 | Forward: CAGCCGTTCTGTCACCGTC |
| | Reverse: CAAGCTGCGCTCAAAGTCC |
| DUSP10 | Forward: TGAAGCACACTCGGATGACC |
| | Reverse: CCTCGAACTCTAGCAACTGCC |
| DUSP16 | Forward: CCTGACTTTATCCCCGAGTCT |
| | Reverse: GAGATCCCAGCTAAACAGTGC |
| MiR-138-5p | Forward: GCGAGCTGGTGTTGTGAATC |
| | Reverse: AGTGCAGGGTCCGAGGTATT |
