## Supplemental Table 2 for "MAPK Pathway Inhibition as A Rational Therapeutic Strategy for MiR-138-5p/PAQR3 Dysregulation-mediated Epirubicin Resistance in Triple-negative Breast Cancer"

**Table S2** Genes in negative regulation of MAPK pathway

| Gene symbol | Annotation |
|---|---|
| PAQR3 | Progestin and adipoQ receptor family member 3 |
| PPP5C | Protein Phosphatase 5 catalytic subunit |
| PPP2R4 | Protein phosphatase 2 phosphatase activator |
| DUSP1 | Dual specificity phosphatase 1 |
| DUSP2 | Dual specificity phosphatase 2 |
| DUSP4 | Dual specificity phosphatase 4 |
| DUSP5 | Dual specificity phosphatase 5 |
| DUSP6 | Dual specificity phosphatase 6 |
| DUSP8 | Dual specificity phosphatase 8 |
| DUSP9 | Dual specificity phosphatase 9 |
| DUSP10 | Dual specificity phosphatase 10 |
| DUSP16 | Dual specificity phosphatase 16 |
