## Supplemental Table 3 for "MAPK Pathway Inhibition as A Rational Therapeutic Strategy for MiR-138-5p/PAQR3 Dysregulation-mediated Epirubicin Resistance in Triple-negative Breast Cancer"

**Table S3. Description of the cohort characteristics (n=114)**

| Characteristics | Number (%) |
|---|---|
| **Age, years** | |
| ≤50 years | 69 (60.5) |
| >50 years | 45 (39.5) |
| **Tumor size** | |
| T1 | 16 (14.0) |
| T2 | 79 (69.3) |
| T3 | 5 (4.4) |
| T4 | 14 (12.3) |
| **Lymph node** | |
| N0 | 52 (45.6) |
| N1 | 57 (50.0) |
| N2 | 3 (2.6) |
| N3 | 2 (1.8) |
| **Histologic grade** | |
| G1 | 2 (1.8) |
| G2 | 20 (17.5) |
| G3 | 87 (76.3) |
| NA | 5 (4.4) |
| **HER2_FISH** | |
| No amplification | 59 (51.8) |
| Amplification | 29 (25.4) |
| NA | 26 (22.8) |
| **TOP2A_FISH** | |
| Deletion | 10 (8.8) |
| Normal | 67 (58.8) |
| Amplification | 10 (8.8) |
| NA | 27 (23.7) |
| **pCR** | |
| No | 98 (86.0) |
| Yes | 16 (14.0) |

pCR, pathologic complete response; FISH, fluorescent in situ hybridization; TOP2A, topoisomerase II-α; HER2, human epidermal growth factor receptor 2.
