## Supplemental Table 4 for "MAPK Pathway Inhibition as A Rational Therapeutic Strategy for MiR-138-5p/PAQR3 Dysregulation-mediated Epirubicin Resistance in Triple-negative Breast Cancer"

### Table S4. SPE, NPV, SEN and PPV in patients receiving epirubicin

| Predictors | Value | 95%CI |
| --- | --- | --- |
| **PAQR3** | | |
| SPE | 0.54 | 0.44-0.64 |
| NPV | 0.93 | 0.83-0.97 |
| SEN | 0.75 | 0.51-0.90 |
| PPV | 0.21 | 0.12-0.33 |
| **TOP2A** | | |
| SPE | 0.50 | 0.40-0.60 |
| NPV | 0.86 | 0.75-0.93 |
| SEN | 0.50 | 0.28-0.72 |
| PPV | 0.14 | 0.07-0.25 |
| **PAQR3_TOP2A** | | |
| SPE | 0.70 | 0.61-0.79 |
| NPV | 0.86 | 0.77-0.92 |
| SEN | 0.31 | 0.14-0.56 |
| PPV | 0.15 | 0.06-0.30 |
| **TOP2A_FISH** | | |
| SPE | 0.95 | 0.87-0.98 |
| NPV | 0.90 | 0.81-0.95 |
| SEN | 0.43 | 0.21-0.67 |
| PPV | 0.60 | 0.31-0.83 |

TOP2A and PAQR3 mRNA binary values were determined at the median of whole cohort. CI, confidence interval; FISH, fluorescent in situ hybridization; TOP2A, topoisomerase II-α; PAQR3, pogestin and adipoQ receptor family member 3; SPE, specificity; NPV, negative predictive value; SEN, sensitivity; PPV, positive predictive value.
