## Supplementary figures and images for "MAPK Pathway Inhibition as A Rational Therapeutic Strategy for MiR-138-5p/PAQR3 Dysregulation-mediated Epirubicin Resistance in Triple-negative Breast Cancer"

### Supplemental Figure 1

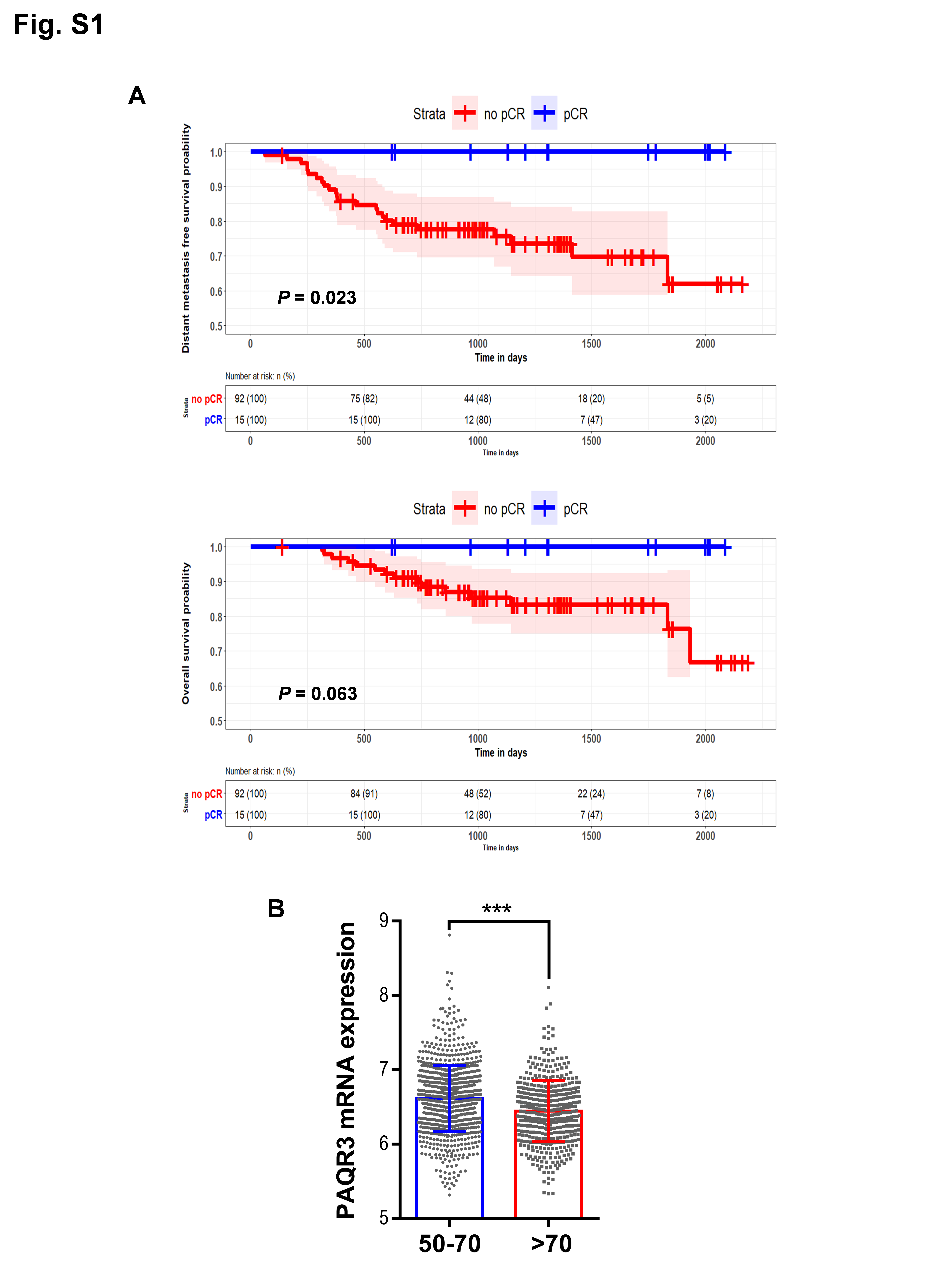

### Supplemental Figure 2

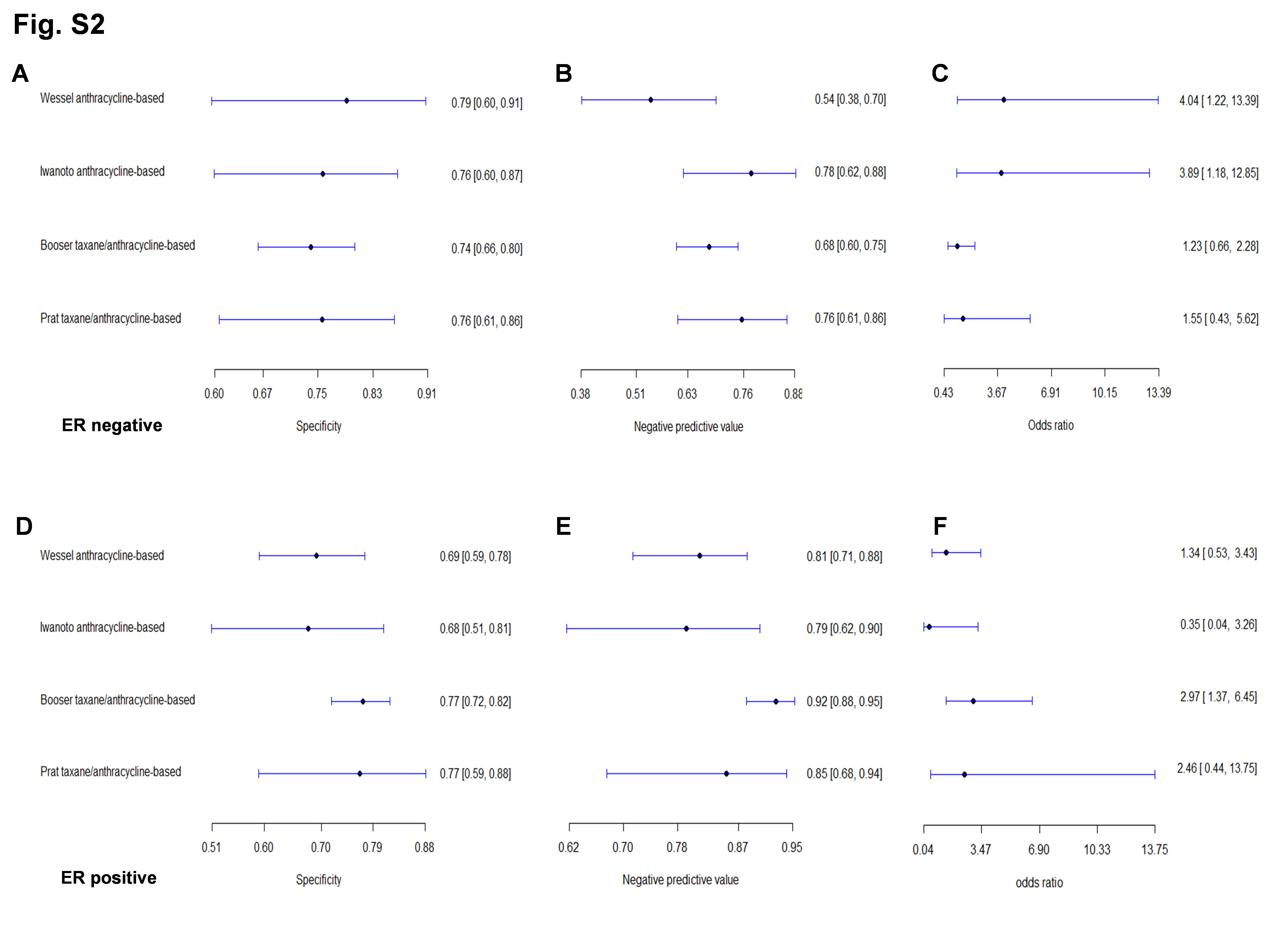
